# Inference of a genome-wide protein-coding gene set of the inshore hagfish *Eptatretus burgeri*

**DOI:** 10.1101/2020.07.24.218818

**Authors:** Kazuaki Yamaguchi, Yuichiro Hara, Kaori Tatsumi, Osamu Nishimura, Jeramiah J. Smith, Mitsutaka Kadota, Shigehiro Kuraku

**Affiliations:** Laboratory for Phyloinformatics, RIKEN Center for Biosystems Dynamics Research, Kobe, Japan; Department of Biology, University of Kentucky, Lexington, KY, USA

## Abstract

The group of hagfishes (Myxiniformes) arose from agnathan (jawless vertebrate) lineages and is one of the only two extant cyclostome taxa, together with lampreys (Petromyzontiformes). Even though whole genome sequencing has been achieved for diverse vertebrate taxa, genome-wide sequence information has been highly limited for cyclostomes. Here we sequenced the genome of the inshore hagfish *Eptatretus burgeri* using DNA extracted from the testis, with a short-read sequencing platform, aiming at reconstructing a high-coverage coding gene catalogue. The obtained genome assembly, scaffolded with mate-pair reads and paired RNA-seq reads, exhibited an N50 scaffold length of 293 Kbp, which allowed the genome-wide prediction of coding genes. This computation resulted in the gene models whose completeness was estimated at the complete coverage of more than 83 % and the partial coverage of more than 93 % by referring to evolutionarily conserved single-copy orthologs. The high contiguity of the assembly and completeness of resulting gene models promises a high utility in various comparative analyses including phylogenomics and phylome exploration.

## Background & Summary

Extant jawless fishes (cyclostomes) are divided into two groups, hagfishes (Myxiniformes) and lampreys (Petromyzontiformes)^1^. They have been studied from various viewpoints mainly because they occupy an irreplaceable phylogenetic position among the extant vertebrates, having diverged from all other vertebrates during the early Cambrian period. Even after massive efforts of whole genome sequencing for invertebrate deuterostomes^2,3^, genome-wide sequence information for species in this irreplaceable taxon was limited until the genome analyses for two lamprey species, the sea lamprey *Petromyzon marinus* and the Arctic lamprey *Lethenteron camtschaticum* were published in 2013 ^4,5^.

In parallel, biological studies involving individual genes have been conducted for both lampreys and hagfishes. Developmental biologists, in particular, have largely relied on lampreys whose embryonic materials are accessible through artificial fertilization^6^, whereas studies on hagfishes have been limited to non-embryonic materials, with a few notable exceptions^7–9^. This type of molecular biological studies is expected to be more thoroughly performed if a comprehensive catalogue of genes is available. For lampreys, derivation of a reliable comprehensive gene catalogue was long hindered by the peculiar nature of protein-coding sequences, which are characterized by high GC-content, codon usage bias, and biased amino acid compositions^5,10,11^. To reinforce existing resources for lampreys, we previously performed a dedicated gene prediction for *L. camtschaticum*^12^ and provided a gene catalogue with comparable or superior completeness to other equivalent resources^4,13^.

As of June 2020, no whole genome sequence information is available for hagfishes except for the one at Ensembl^13^ that remains unpublished, a fact that hinders the comprehensive characterization of gene repertoires and their expression patterns. Currently, some efforts for genome sequencing and analysis are ongoing that aim to resolve large-scale evolutionary and epigenomic signatures^9^, inspired partly by the relevance of hagfish to understanding patterns of whole genome duplications^14–19^ and chromosome elimination^20–22^. In contrast to those efforts, which are necessarily targeting reconstruction of the genome at chromosome scale, in this study we aimed at providing a data set covering as many full-length protein-coding genes as possible, to enable gene-level analysis on molecular function and evolution of hagfishes, an indispensable component of the vertebrate diversity.

## Methods

### Genome sequencing

Genomic DNA was extracted from the testis of a 48cm-long male individual of *Eptatretus burgeri* caught at the Misaki Marine Station in June 2013, with phenol/chloroform as previously described^23^, and the genome sequencing was performed as outlined in Figure 1. The study was conducted in accordance with the institutional guideline Regulations for the Animal Experiments by the Institutional Animal Care and Use Committee (IACUC) of the RIKEN Kobe Branch. The extracted genomic DNA was sheared with an S220 Focused-ultrasonicator (Covaris) to retrieve DNA fragments of variable length distributions (see Table 1 for detailed amounts of starting DNA and conditions for shearing). The sheared DNA was used for paired-end library preparation with a KAPA LTP Library Preparation Kit (KAPA Biosystems). The optimal numbers of PCR cycles for individual libraries were determined with a Real-Time Library Amplification Kit (KAPA Biosystems) by preliminary qPCR-based quantification using an aliquot of adaptor-ligated DNAs as described previously^24^. Small molecules in the prepared libraries were removed by size selection using Agencourt AMPure XP (Beckman Coulter). The numbers of PCR cycles and conditions of size selection for individual libraries are included in Table 1. Mate-pair libraries were prepared using a Nextera Mate Pair Sample Prep Kit (Illumina), employing our customized iMate protocol^25^ (http://www.clst.riken.jp/phylo/imate.html). The detailed conditions of mate-pair library preparation are included in Table 1. After size selection, the quantification of the prepared libraries was performed using a KAPA Library Quantification Kit (KAPA Biosystems). They were sequenced on a HiSeq 1500 (Illumina) operated by HiSeq Control Software v2.0.12.0 using a HiSeq SR Rapid Cluster Kit v2 (Illumina) and HiSeq Rapid SBS Kit v2 (Illumina), HiSeq X (Illumina) operated by HiSeq Control Software v3.3.76, and MiSeq operated by MiSeq Control Software v2.3.0.3 using MiSeq Reagent Kit v3 (600 Cycles) (Illumina). Read lengths were 127 or 251?nt on HiSeq 1500, 151 nt on HiSeq X, and 251Cnt on MiSeq. Base calling was performed with RTA v1.17.21.3, and the fastq files were generated by bcl2fastq v1.8.4 (Illumina) for HiSeq 1500 and MiSeq, while RTA v2.7.6 and bcl2fastq v2.15.0 (Illumina) were instead employed for HiSeq X. Removal of low-quality bases from paired-end reads was processed by TrimGalore v0.3.3 (https://www.bioinformatics.babraham.ac.uk/projects/trim_galore/) with the options ‘-- stringency 2 --quality 30 --length 25 --paired --retain_unpaired’. Mate-pair reads were processed by NextClip v1.1 ^26^ with the default parameters.

**Table 1.** Properties of prepared sequencing libraries.

A. Paired-end genome shotgun libraries
| Accession ID | Library ID | Average insert size (bp) | Amount of DNA used (µg) | # PCR cycles | # read pairs |
| --- | --- | --- | --- | --- | --- |
| DRX218807,<br>DRX218808,<br>DRX218809 | P167_02_1 | 420 | 0.05 | 3 | 185,747,472 |
| DRX218810,<br>DRX218811,<br>DRX218812,<br>DRX218813 | P167_02_2 | 690 | 0.05 | 5 | 174,756,277 |
| DRX218814,<br>DRX218815 | P167_12_1 | 644 | 3 | 0 | 127,124,057 |
| DRX218816,<br>DRX218817,<br>DRX218818 | P167_12_2 | 873 | 3 | 4 | 278,906,224 |
| DRX218819,<br>DRX218820,<br>DRX218821 | P167_12_4 | 381 | 3 | 0 | 329,285,268 |
| DRX218822,<br>DRX218823,<br>DRX218824 | P167_12_5 | 418 | 3 | 2 | 129,764,303 |

B. Mate-pair genome libraries
| Accession ID | Library ID | Mate distance (Kb) | Amount of DNA used (µg) | # PCR cycles | # read pairs |
| --- | --- | --- | --- | --- | --- |
| DRX218825 | P167_02_5 | 6-10 | 4 | 10 | 9,632,719 |
| DRX218826 | P167_02_6 | 12-18 | 4 | 13 | 10,230,697 |
| DRX218827,<br>DRX218828 | P167_02_7 | 6-10 | 4 | 10 | 219,246,424 |
| DRX218829,<br>DRX218830 | P167_02_8 | 12-18 | 4 | 13 | 139,814,746 |

C. RNA-seq libraries
| Accession ID | Library ID | Tissue | Amount of total RNA used (µg) | # PCR cycles | # read pairs |
| --- | --- | --- | --- | --- | --- |
| DRX218831 | P238_01_1 | liver | 1 | 6 | 55,675,220 |
| DRX218832 | P238_02_1 | blood | 1 | 5 | 57,600,815 |

**Figure 1.**
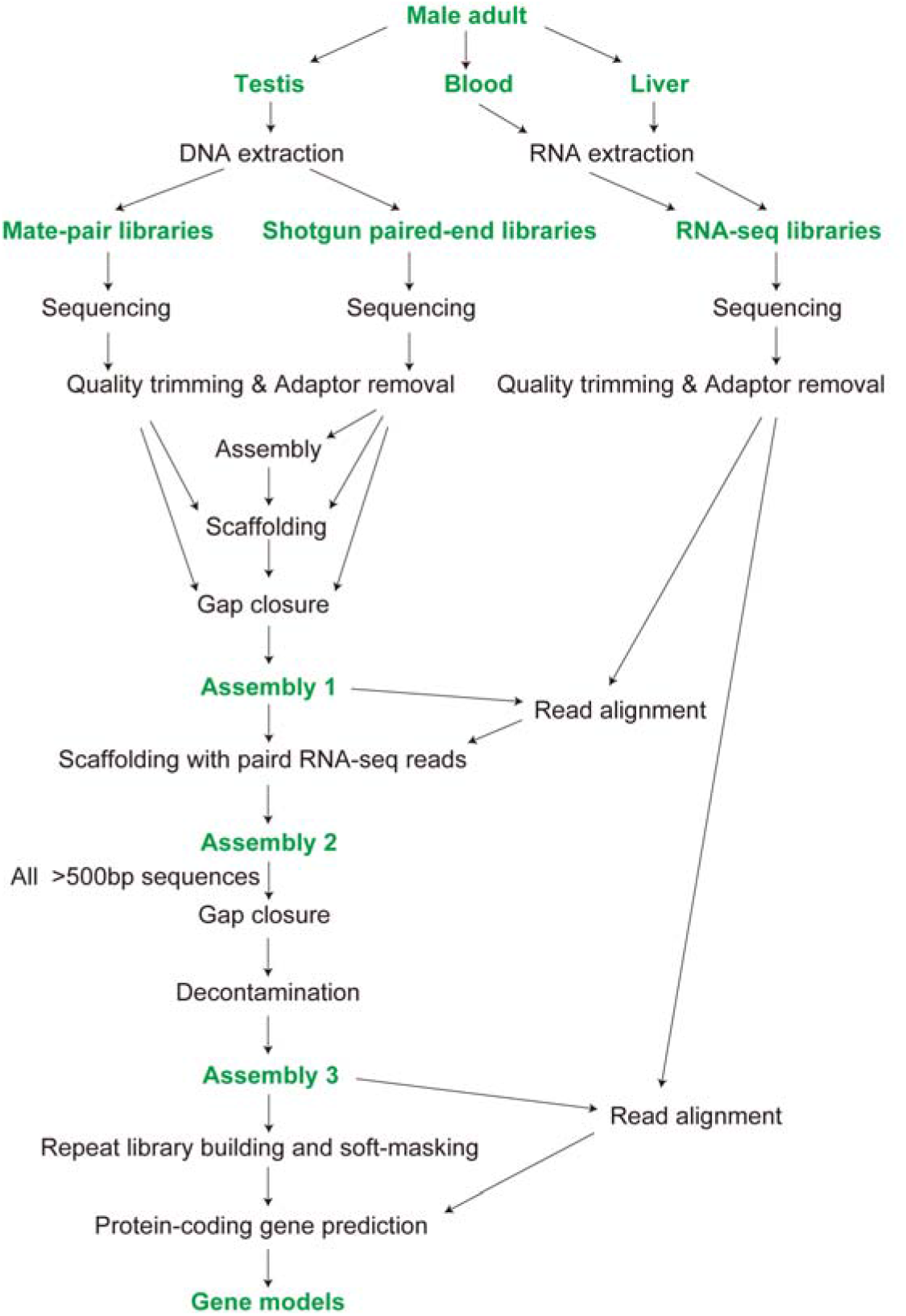
Data production workflow. Samples, raw data, and products are indicated with green letters, while computational steps are labelled in black. See Methods for the details including the choice of the programs used in individual computational steps.

**Figure 2.**
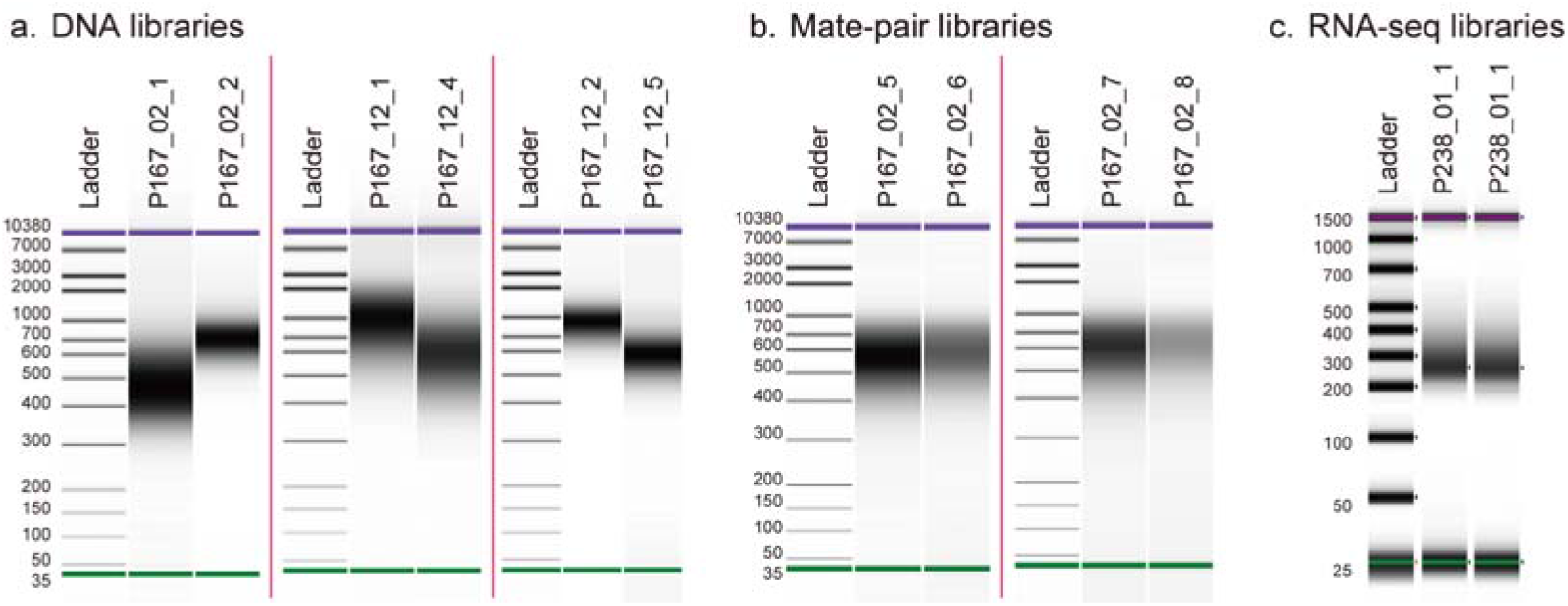
Length distributions of DNA molecules in sequencing libraries. a, Shotgun DNA libraries. b, Mate-pair libraries. c, RNA-seq libraries.

### RNA-seq and transcriptome data processing

Total RNAs were extracted from the liver and the blood of the above-mentioned adult individual with Trizol reagent (Thermo Fisher Scientific). Quality control of DNase I-treated RNA was performed with Bioanalyzer 2100 (Agilent Technologies), which yielded the RIN values of 8.7 and 9.1 for the respective tissues. Libraries were prepared with TruSeq Stranded mRNA LT Sample Prep Kit (Illumina) as previously described^27^. The amount of starting total RNA and numbers of PCR cycles are included in Table 1. The obtained sequence reads were trimmed for removal of adaptor sequences and low-quality bases with TrimGalore v0.3.3 as outlined above. Alignment of the trimmed RNA-seq reads to the genome assembly employed HISAT2 v2.2.1 ^28^ with the options of ‘-k 3 -p 20 --pen-noncansplice 1000000’.

### Genome assembly

*De novo* genome assembly and scaffolding that employs the processed short reads were carried out by the program PLATANUS v1.2.4 ^29^ with its default parameters. The assembly step employed paired-end reads and single reads whose pairs had been removed, and the scaffolding step employed paired-end and mate-pair reads. The gap closure step employed all of the single, paired-end and mate-pair reads after processing. The obtained sequences were further scaffolded with paired-end RNA-seq reads with the program P_RNA_Scaffolder^30^ (commit 7941e0f on May 30, 2019, at GitHub) with the options ‘-s yes -b yes -p 0.90 -t 20 -e 100000 -n 100’, followed by another gap closure run with PLATANUS ‘gap_closure’ using the same set of reads used in the above-mentioned gap closure run. Resultant genomic scaffold sequences were screened for own mitochondrial DNA fragments, contaminating organismal sequences, PhiX sequences loaded as a control, mitochondrial DNA sequences, and those shorter than 500⍰bp, as performed previously^31^.

### Repeat detection and masking

To obtain species-specific repeat libraries, RepeatModeler v1.0.8 ^32^ was run on the genome assemblies of the individual species with default parameters. Detection of repeat elements in the genomes was performed by RepeatMasker v4.0.5 ^33^, which employs the National Center for Biotechnology Information (NCBI) RMBlast v2.2.27 ^34^, using the custom repeat library obtained above. For gene prediction, the parts of genome sequences detected as repeats are soft-masked by RepeatMasker with the options ‘-nolow -xsmall’.

### Construction of gene models

Construction of gene models was performed by employing the gene prediction pipeline Braker v2.1.4 ^35^ with the options ‘--min_contig=500 --prg=gth --softmasking --UTR=off’ (Figure 1). This computation employed RNA-seq read alignments in a.bam file onto the genome assembly and a set of peptide sequences prepared as following. The set of peptide sequences used as homolog hints included the predicted proteins of the Arctic lamprey (34,362 sequences, previously designated as GRAS-LJ^12^), which were aligned to the soft-masked genome assembly.

## Data Records

### Genome assembly

Our technical procedure employing the genome assembly program PLATANUS^29^ that previously produced genome assemblies for multiple shark species with modest investment^36^ yielded genome sequences consisting of 4,519,897 scaffolds (Assembly 1 in Figure 1) with an N50 length of 238 Kbp (length cutoff = 500 bp). To improve the continuity of fragmentary sequences that were derived from transcribed regions but were separated from exons, the sequences in Assembly 1 were further scaffolded with paired-end RNA-seq reads, which resulted in 4,505,643 sequences (Assembly 2) with an N50 length of 264 Kbp (length cutoff = 500 bp). These sequences were filtered for the length of > 500 bp, processed again for gap closure with the program PLATANUS, and scanned for contaminants of microbes and artificial oligos used for sequencing. Through this procedure, we have obtained 114,941 sequences with the minimum and maximum lengths of 500bp and 2.064 Mbp, respectively, marking the N50 scaffolding length of 293 Kbp (Assembly 3). These sequences have systematic identifiers scf_eptbu00000001□scf_eptbu00114941 and are available under https://figshare.com/projects/eburgeri-genome/77052.

### Gene models

Using the resultant genome sequences (Assembly 3), genome-wide prediction of protein-coding sequences were performed with the program pipeline Braker^35^. After preliminary runs with variable parameters and input data sets, we conducted a prediction run with transcript evidence and peptide hints, which resulted in a set of 46,295 genes, with the maximum length of the putative peptides of 19,580 amino acids. These sequences have systematic identifiers Eptbu0000001□Eptbu0046295 with suffixes ‘.t1’□‘.t6’ depending on the multiplicity of predicted peptide variants derived from alternative splicing. These sequences are available under https://figshare.com/projects/eburgeri-genome/77052.

## Technical Validation

### Mapping RNA-seq reads to the genome assembly

To confirm the coverage of the genome assembly, paired-end RNA-seq reads were aligned with splicing-aware read mapping program HISAT2 as described in Methods. This computation resulted in the high percentage of the paired reads mapped to the nuclear genome sequences of 91.64 % while the majority of the remaining reads (5.17%) were mapped to the mitochondrial genome sequence of *E. burgeri* itself.

### Gene space completeness assessment of genome assembly and gene models

It has been previously shown that completeness scores of cyclostome genomes tend to be underestimated, when their rapid-evolving nature and phylogenetic position is not taken into consideration^27^. In this study, completeness of the genome assemblies was assessed with CEGMA v2.5 ^37^ and BUSCO v2.0.1 ^38^. For both CEGMA and BUSCO, we employed not only the reference gene sets provided with these pipelines but also the core vertebrate genes (CVG) that was developed specifically for vertebrates from isolated lineages such as elasmobranchs and cyclostomes^27^. The assessments were executed on the gVolante webserver^39,40^. The percentages of single-copy orthologs detected as ‘complete’ was approximately 65 %, with 91 % detected as either ‘complete’ or ‘partial/fragmented’, when CEGMA was used with CVG.

Similarly, we performed completeness of the gene models with BUSCO v2.0.1 ^38^, again using CVG as a reference ortholog set. The assessment resulted in the complete coverage of 83.7 % and partial coverage of 93.6 % of CVGs (Table 2). The difference of the completeness scores between the assessments of the genome assembly and the gene models might be explained by decreased sensitivity of detecting divergent multi-exon genes in the genome. Altogether, the resultant set of gene models is expected to encompass more than 90 % of the protein-coding genes in the *E. burgeri* genome.

**Table 2.** Statistics of the newly produced gene models compared with published cyclostome gene models.

| Species | Source | # Genes<br>(# Peptides) | Maximum<br>peptide length<br>(amino acids) | Completeness score<sup>b</sup> (%) |  |
| --- | --- | --- | --- | --- | --- |
|  |  |  |  | Only<br>'Complete' | Including<br>'Fragmented' |
| *Eptatretus burgeri* | This study | 46295<br>(50127) | 19580 | 83.7 | 93.6 |
| *Lentheteron camtschaticum* | GRAS-LJ<sup>12,a</sup> | 34435 | 19612 | 90.1 | 98.7 |
| *Petromyzon marinus* | PMZ_v3.0<sup>19</sup> | 20940<br>(20950) | 18818 | 57.1 | 89.3 |
| *Petromyzon marinus* | Ensembl<br>gene build<sup>13</sup> | 10415<br>(11442) | 18900 | 84.1 | 94.9 |
| *Petromyzon marinus* | PMZ1.0<sup>5</sup> | 24132<br>(24271) | 17467 | 63.5 | 89.3 |

<sup>a</sup>The construction of this gene model was performed without predicting alternative splice variants, and the number of peptides is thus not included in the relevant cell.
<sup>b</sup>The completeness was scored by the use of the pipeline BUSCO v2 with the one-to-one ortholog set CVG (see Methods).

## Data Citations

The sequence data are available in fastq files at DDBJ DRA under the accession ID DRA010216 and as multifasta files at https://figshare.com/projects/eburgeri-genome/77052, as well as formatted databases for BLAST searches at https://transcriptome.riken.jp/squalomix/.

## Usage Notes

This data set is oriented towards gene-level analysis including phylogenomic analysis and phylome exploration aiming at studying gene family evolution, rather than the analysis of complete genome structure. Importantly, the total length of the genome sequences obtained in this study amounts only to approximately 1.7 Gbp which is smaller by more than 1 Gbp than the genome size estimate based on flow cytometry of nuclear DNA content^21^ (2.91 Gbp). For investigating the structural evolution of the whole genome, such as chromosome elimination or large-scale synteny conservation, it may be advisable to wait for other resources to be released without embargo.

The sequence files of the obtained gene models sometimes include multiple transcripts and its deduced amino acid sequences per gene, because of predicted alternative splice variants. To facilitate the use of the dataset without splice variants, a sequence file without splice variants (doi: 10.6084/m9.figshare.11971932) has also been made available.

## Code Availability

No custom computer code was employed in this study.

## Acknowledgements

We thank Masumi Nozaki for assistance in sampling. The authors acknowledge Kazu Tanimoto, Kaori Tanaka, and Chiharu Tanegashima at Laboratory for Phyloinformatics in RIKEN Center for Biosystems Dynamics Research (BDR) for technical assistance. This study was supported by RIKEN and JSPS KAKENHI Grant Number 17K07426 to S.K. and an NSF Grant Number MCB-1818012 to J.J.S‥

## Author contributions

S.K conceived the study. J.J.S. sampled the tissues. Y.H., K.Y., O.N., M.K, and S.K performed experiments. Y.H., K.Y., J.J.S. and S.K interpreted data. S.K drafted the manuscript. All authors contributed to the final manuscript editing.

